# A simple and sensitive SYBR Gold-based assay to quantify DNA-protein interactions

**DOI:** 10.1101/634816

**Authors:** Spencer Schreier, Bhanu Prakash Petla, Tao Lin, Suvobrata Chakravarty, Senthil Subramanian

## Abstract

A simple, accessible, and inexpensive assay to quantify the strength of DNA-protein interactions was developed. The assay relies on capturing DNA-protein complexes using an affinity resin that binds tagged, recombinant proteins. Sequential washes with filtration spin cups and centrifugation remove non-specific interactions in a gentle, uniform manner and a final elution isolates specific DNA-protein complexes. SYBR Gold nucleic acid stain is added to the eluted product and the fluorescence intensity accurately quantifies the amount of captured DNA, ultimately illustrating the relative strength of the DNA-protein interaction. The major utility of the assay resides in the versatility and quantitative nature of the SYBR Gold:nucleic acid interaction, eliminating the need for customized or labeled oligos and permitting relatively inexpensive quantification of binding capacity. The assay also employs DNA-protein complex capture by the very common purification tag, 6xHis, but other tags could likely be utilized. Further, SYBR Gold fluorescence is compatible with a wide variety of instruments, including UV transilluminators, a staple to any molecular biology laboratory. This assay was used to compare the binding capacities of different Auxin Response Factor (ARF) transcription factors to various dsDNA targets, including the classical AuxRE motif and several divergent sequences. Results from dose-response assays suggest that different ARF proteins might show distinct comparative affinities for AuxRE variants, emphasizing that specific ARF-AuxRE binding strengths likely contribute to the complex and fine-tuned cellular auxin response.

## Introduction

The relationship between DNA-binding proteins and their genomic targets is fundamental to an organism’s health, development, and adaptivity. Several assays have been developed to study such important interactions, such as the Electrophoretic Mobility Shift Assay (EMSA) (Hellman and Fried 2007), ChIP-Seq (Robertson *et al.* 2007), high-throughput protein binding microarrays (Berger and Bulyk 2009), and surface plasmon resonance (Wegner *et al.* 2003). While powerful, these methods collectively have drawbacks concerning timeliness, throughput, accuracy, and experimental feasibility. A DNA-protein binding assay that relies on common molecular biology tools and supplies would have great utility for research laboratories, teaching laboratories or research groups that do not have access to the sophisticated equipment, time commitment, or expensive resources that are characteristic of the aforementioned assays.

Several plant hormones manifest their activity through specific protein-DNA interactions, such as a transcription factor binding to specific regulatory elements in the DNA. Auxin is a critical phytohormone whose activity contributes to nearly every aspect of plant growth and development, yet still needs further understanding. The regulatory mechanism underpinning auxin activity is a dynamic relationship between receptors (Tan *et al.* 2007), transcriptional activators and repressors (Tiwari *et al.* 2003), as well as epigenetic players (Nizampatnam *et al.* 2015, Wu *et al.* 2015). The activator auxin response factors (ARFs) reside at the core of the auxin signaling pathway and have been well characterized. Activator ARFs engage auxin-responsive gene promoters, but are withheld from inducing an unlicensed auxin response in the absence or under low concentrations of auxin by the Aux/IAA repressor proteins (Han *et al.* 2014). When auxin complexes with the TRANSPORT INHIBITOR RESPONSE 1 (TIR1) receptor and stabilizes the assembly of an E3 ubiquitin ligase complex with the S-Phase Kinase Associated Protein 1-Cullin-F-Box Protein (SCF) (Dharmasiri *et al.* 2005, Tan, et al. 2007), the Aux/IAA repressors are degraded to allow the activator ARF to induce auxin-responsive genes. Distinct interaction patterns between different ARFs and Aux/IAA proteins govern part of the complexity and specificity in auxin responses (Piya *et al.* 2014, Weijers and Wagner 2016).

Activator ARFs have been long known to bind a TGTCTC DNA motif (Ulmasov *et al.* 1995) proximal to auxin-responsive genes, yet more recent discoveries into the activity of the ARF proteins themselves reveal nuances in sequence specificity and motif conformation (Boer *et al.* 2014). For example, dimerization within the ARF DNA binding domain led to improved affinity for auxin response elements (AuxREs) and the palindromic nature of the AuxRE motif coincides with this activity. Importantly, ARFs evaluated in the above study showed a preference to a TGTCGG motif over the classical TGTCTC AuxRE motif. In addition to sequence specificity, the spacing between AuxRE motifs also contributed to binding strength. This sponsors a key question: do the affinities of each ARF protein vary among the different AuxREs? A family of ARFs with preferential binding to specific AuxRE motifs would indeed allow auxin to dictate complex transcription responses, enabling regulation of multiple plant processes.

We sought to develop a flexible and feasible binding assay that would allow us to test different AuxRE variants against ARF proteins. We validated the assay using Arabidopsis ARF5 (AtARF5) and soybean ARF8a (GmARF8a). Binding preference of AtARF5 was studied in detail recently using surface plasmon resonance and protein-binding microarrays (Boer, et al. 2014). GmARF8a regulates auxin signaling during symbiotic nodule development, a legume-specific developmental process (Wang *et al.* 2015). We show that individual activator ARFs have their own specific affinity to various AuxRE sequences. Furthermore, we illustrate the importance of a palindromic sequence conformation within the AuxRE for stable ARF affinity. Beyond the scope of these findings, we also demonstrate that this assay is sensitive, timely, and operable using staple molecular biology equipment. Furthermore, the assay has likely utility for studying other types of molecular interactions in a timely, flexible manner.

## Materials and Methods

### ARF DBD Selection, Design, and Cloning

The DNA binding domain (DBD) of two ARF proteins classified as activator ARFs were selected to compare their relative DNA binding capacity. The peptide sequence of *Glycine max* ARF8a (Glyma02g239600.1, genome version Glyma2.0; GmARF8a) was aligned to the 23 *Arabidopsis thaliana* ARFs (Clustal Omega, EMBL-EBI) to identify domain boundaries. Previously, Boer et al. described the mechanistic action of the *A. thaliana* ARF5 and ARF1 DBDs (AtARF5 and AtARF1 respectively; Boer, et al. 2014), and these same ARF domains served as a selection scaffold for the GmARF8a DBD. Boer et al. expressed the DNA binding domains of AtARF1 and AtARF5 proteins, incorporating residues 1-354 and 1-390, respectively. Based on this range, primers were designed to clone residues 1-362 for GmARF8a, along with the same AtARF5 DBD studied by Boer et al. (Table S2). These primer sequences also added C-terminal 6× histidine tags for downstream purification and application in the DNA binding assay described herewith.

Each ARF DBD was cloned in the pETite N-His SUMO Kan expression vector (Lucigen, WI) according to the manufacturer’s instructions. Verified constructs were transformed into Rosetta2 DE3 expression strain (EMD Millipore, Germany) and were selected on kanamycin/chloramphenicol plates (50 and 20 μg/ml, respectively).

### Protein Purification

Clones harboring each ARF were cultured in LB/kanamycin/chloramphenicol media to an O.D_600_ of 0.75 and induced for expression with 0.5mM IPTG (Gold Biotechnology, MO) overnight at 16°C. Buffers were prepared per Table S3, filter sterilized, and stored at 4°C until use. All purification steps were performed at 4°C. The cells were harvested by centrifugation, resuspended in lysis buffer, and lysed by sonication. The lysed cells were centrifuged and the supernatant filtered and applied to a single 1ml His-Trap FF (GE Healthcare Life Sciences, WA) column that was dedicated to each ARF. The column was washed with 40ml wash buffer and the ARFs eluted with 250mM elution buffer in 1ml fractions. Individual fractions were run on SDS PAGE to check for purity. The pETite N-His SUMO Kan expression vector that these ARFs were cloned into generates a SUMO tag at the N-terminus of the proteins of interest which facilitates better bacterial expression. To ensure that this tag would not interfere with ARF:AuxRE interactions, each purified recombinant ARF was treated with SUMO Express Protease (Lucigen, WI) at 4°C overnight to cleave this tag. Fractions corresponding to the molecular weight of the native ARFs were confirmed for purity using SDS PAGE and pooled together. The pooled fractions were exchanged into SPR buffer (Table S3) using a 20ml Pierce Concentrator with a 10,000 molecular weight cut-off (ThermoFisher, CA) to remove the SUMO tag. The exchanged sample was diluted to either 0.8 or 0.4mg/ml via the Bio-Rad Protein Assay Dye Reagent Concentrate (Bio-Rad, CA). Samples were divided into 100μL aliquots and flash frozen with liquid nitrogen. The samples were immediately stored at −80°C until use.

### Dose-responsive Binding Assay Details

To resolve ARF-DNA interactions, a centrifugation-based binding assay was developed. This assay is summarized in Figure 1. Prior to the assay, Auxin Response Element (AuxRE) sequences mimicking the binding sequences tested by Boer et al. (Boer, et al. 2014), as well as divergent sequences, were ordered as single-stranded oligos (Table S1). These oligos were resuspended in Type 1 water to 100μM. Each double-stranded AuxRE target was generated by mixing equal volumes (50:50μM) of the AuxRE sequence complements, placing in a beaker filled with 95°C water and allowing to cool to room temperature for annealing. The ARF, AuxRE target, and SPR buffer were combined in a small volume (20-50μ L) binding reaction in a 2ml, round-bottom microfuge tube (Eppendorf) and placed on a nutator set to 24 RPM in a 4°C cold room for 1 hour and 15min. Three independent binding reactions were performed for each assay. It is important to use a round-bottom microfuge tube over a conical type, as it allows better inversion and mixing for the assay. A slurry containing 5μL of GE High Performance Ni-NTA resin (GE Healthcare Life Sciences, PA) and 175μL of SPR buffer was added to each tube and continued to incubate at 24 RPM for 15min. The tube was centrifuged at 100 × g for 30 sec to pull down any slurry from the lid and 300μ L of SPR buffer was added and mixed. The entire 500μ L sample was transferred to a Pierce Spin Cup (ThermoFisher, CA) and centrifuged at 500 × g for 2min at 4°C in a swinging bucket rotor. The flow through was discarded and 500μ L of SPR buffer was added to the column for further washing. The column was centrifuged at 500 x g for 2 min and transferred to a fresh 2ml microfuge tube. 110μ L of 0.5M EB (Table S3) was added to the column and allowed to incubate for 10min. The column was centrifuged as before and 100μ L of the flow through was transferred to a well of a black Corning microplate (ThermoFisher, CA) containing 1μ L of 100x SYBR Gold Nucleic Acid Stain (ThermoFisher, CA). A reaction where no protein (ARF) was added was used as control. The plate was incubated on a shaker set to 200 RPM for 15 min at room temperature. Fluorescence was measured using a Biotek Synergy 2 plate reader (Biotek, VT) at 485/520 excitation and emission at level 75 sensitivity.

**Figure 1.**
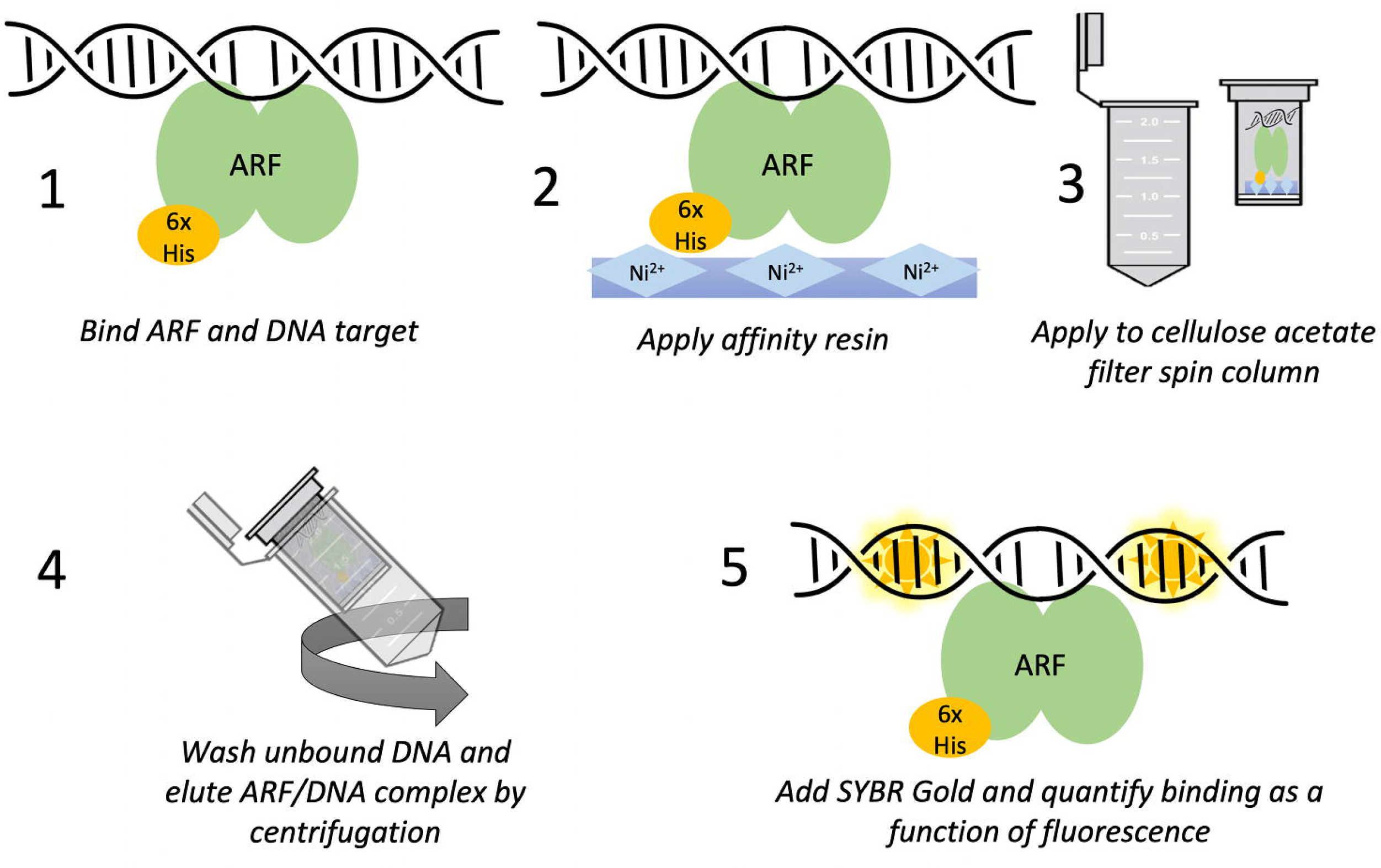
Steps involved in SIPDI assay used to evaluate and quantify ARF binding to the DNA targets tested in this study.

To evaluate if the assay is accessible to commonly available lab equipments, the plate was also imaged on a UVP High Performance UV Transilluminator and LI-COR Odyssey Fc (LI-COR, NE). For the UV transilluminator measurement, the plate was exposed for 0.386 min on low UV intensity and captured with a Kodak Gel Logic 112 camera system. The plate image was saved as a .tiff file and analyzed in ImageJ by selecting each well as a region of interest and its mean fluorescence collected (Figure S2). For the LI-COR Odyessy Fc, the plate was exposed for 2 min with the 600 nm channel and the second-brightest image selected for analysis. The fluorescence intensity of each well was measured using Image Studio software and delivered as a total fluorescence value (Figure S3). In all cases, ANOVA tests were carried out between samples with an α=0.05 (Julkowska *et al.* 2019).

## Results

### Principle of the <u>S</u>YBR Gold-based <u>i</u>n vitro <u>p</u>rotein-<u>D</u>NA <u>i</u>nteraction (SIPDI) assay

The assay involves (i) combining the purified protein and dsDNA molecules of interest in an optimal binding environment (Step 1; Figure 1), (ii) pulling down DNA-protein complexes from the mixture using an affinity tag (Steps 2, 3, and 4; Figure 1), and (iii) quantifying the amount of bound dsDNA using SYBR Gold fluorescence (Step 5; Figure 1).

Using His-tagged proteins and Ni-NTA resin allowed us to vary the amount of protein used as needed, enabling quantitative estimates of binding affinity. The higher sensitivity of the fluorescent plate reader to detect SYBR Gold allowed the use of unlabeled dsDNA in the assay, making it inexpensive to test various dsDNA targets (see Experimental Procedures for details).

### SIPDI assay closely reproduced protein microarray results for AtARF5-auxRE interaction

It was previously shown using a protein-binding microarray that AtARF5-DBD bound with different affinities to variants of the AuxRE. For example, highest preferential binding was observed with the TGTCGG motif while TGTCAT showed much lower affinity to AtARF5-DBD (Boer, et al. 2014). We evaluated the ability of a SYBR Gold-based assay to reproduce these differences. Five different variants of AuxRE dsDNA (Table S1) were synthesized and used in the assay. The observed binding capacities measured by SIPDI assay (Figure 2) closely reflected the results from protein-binding microarray experiments by Boer, et al. (2014). For example, ER7-AT had the lowest binding to AtARF5-DBD, followed by ER7-GA and ER5. The canonical AuxRE ER7 had significantly higher binding to AtARF5-DBD vs. the above AuxREs while ER7-GG had the highest binding to AtARF5-DBD.

**Figure 2.**
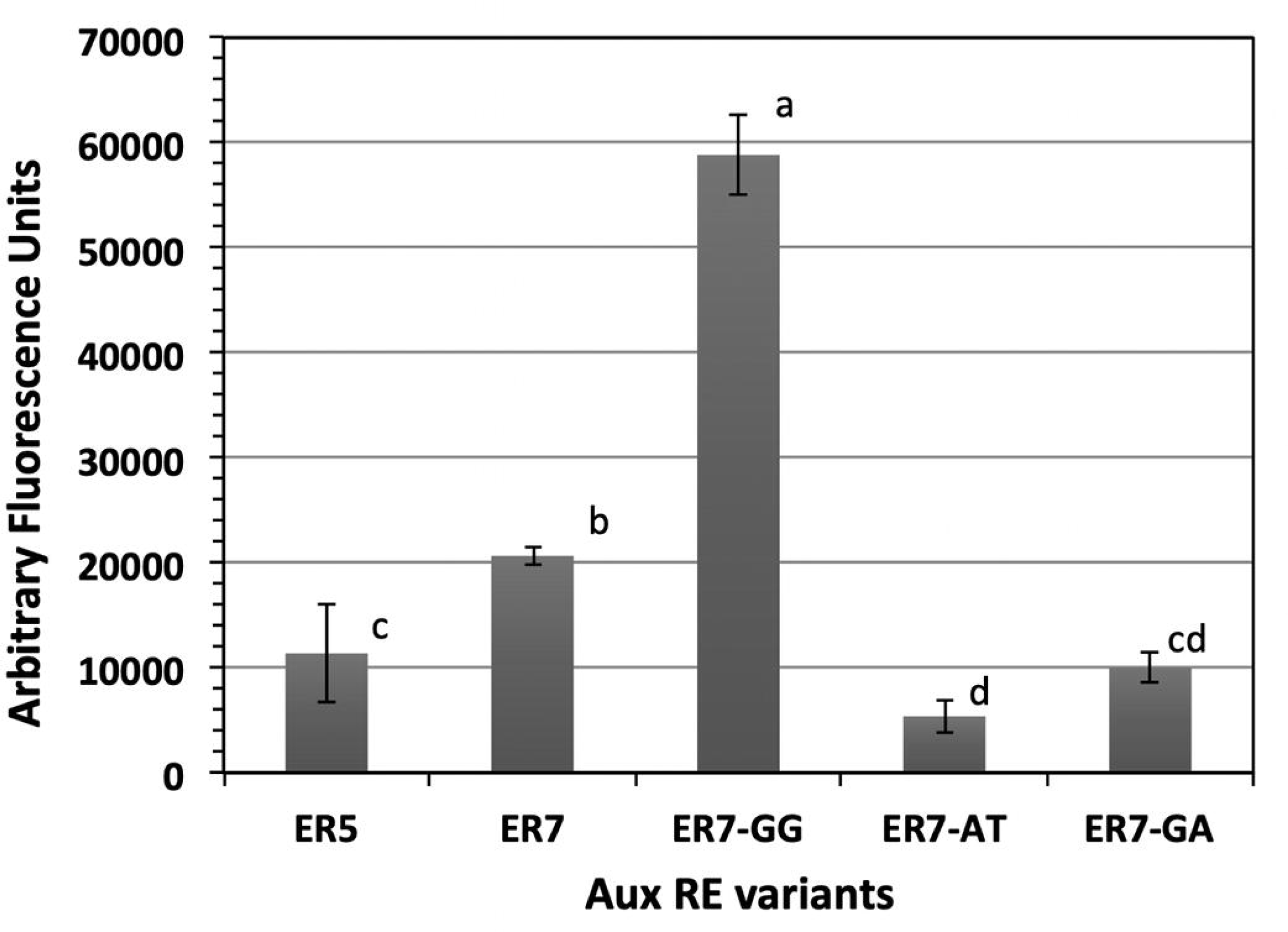
Binding of AtARF5 to different AuxRE variants. Amount of AuxRE dsDNA co-eluted with AtARF5 was measured by SYBR-Gold fluorescence. Data shown are average of three independent binding assays after subtracting fluorescence in no protein controls for each AuxRE. Error bars indicate standard deviation. Fluorescence measurements of samples labeled with distinct letters are significantly different from each other (ANOVA test, P < 0.05).

Therefore, we concluded that the SYBR Gold-based assay can be used to quantify relative affinities of different protein-DNA interactions.

### SIPDI assay has a wide dynamic range

We evaluated whether the SYBR Gold nucleic acid stain generates fluorescence in a linear manner with increasing concentrations of AuxREs. A concentration range of 0.01-0.06 μM indeed produced a linear fluorescence output with SYBR Gold (Figure S1). To evaluate the dynamic range of the assay for ARF-AuxRE binding, we tested the binding of 2, 4, 6, and 8 μM of ER7-GG, (the AuxRE to which AtARF5-DBD bound with highest affinity) against 4μg of AtARF5-DBD (4.44 μM) and GmARF8a-DBD (5.0 μM). Increase in binding was observed with increasing quantities of ER7-GG up to 4 μM for GmARF8a-DBD, and up to 6 μM for AtARF5-DBD (Figure 3). This indicated that 6 μM of the AuxRE target dsDNA was not rate-limiting for the binding assay.

**Figure 3.**
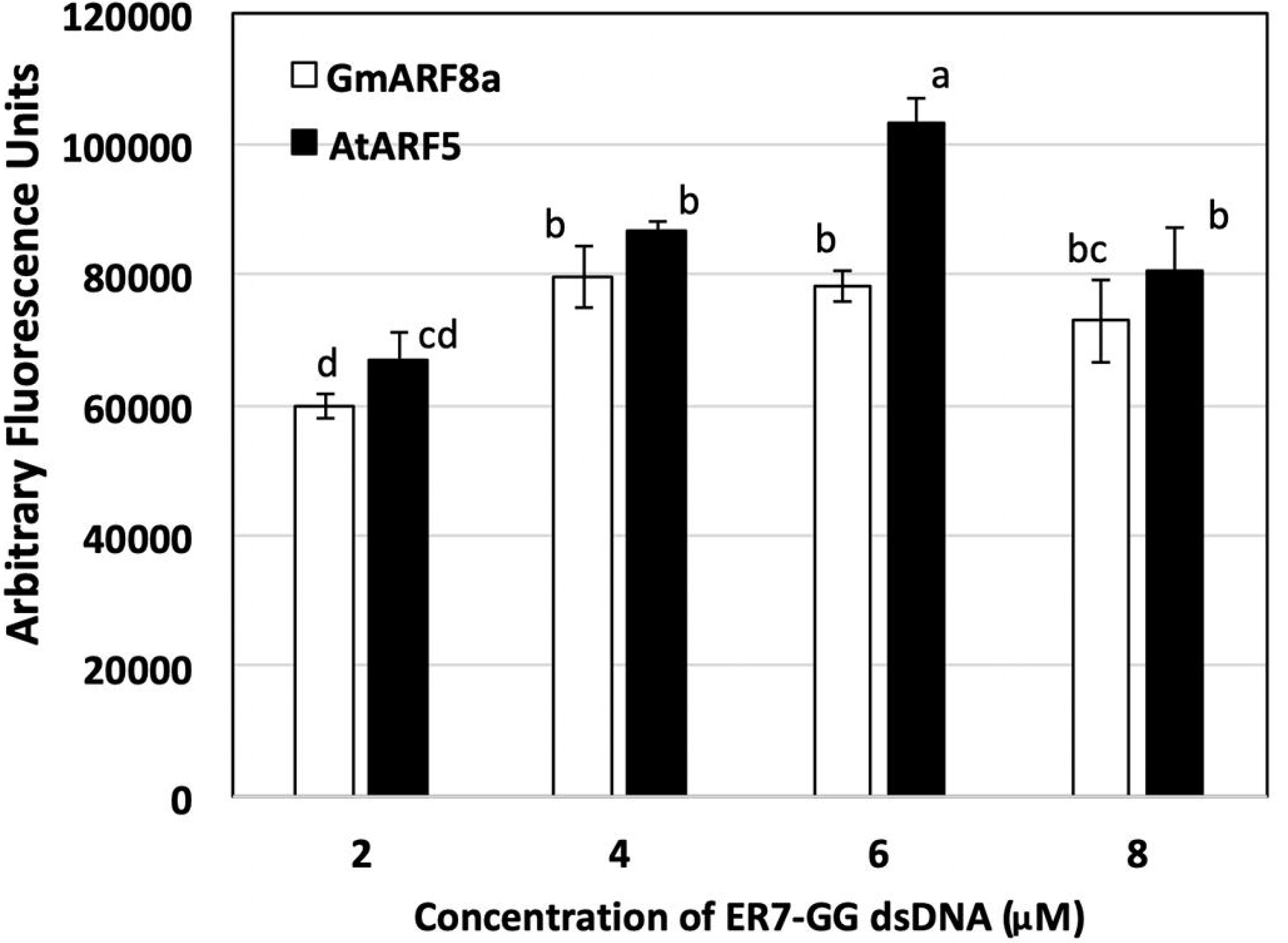
Binding of AtARF5 and GmARF8a to increasing amounts of ER7-GG. Data shown are average of three independent binding assays after subtracting fluorescence in no protein controls for each AuxRE. Error bars indicate standard deviation. Fluorescence measurements of samples labeled with distinct letters are significantly different from each other (ANOVA test, P < 0.05)

Next, we evaluated binding when increasing amounts of ARF-DBDs were combined with ER7-GG and the canonical AuxRE ER7 (TGTCTC). As 6μM of target appeared to be the saturating concentration (Figure 3), 6μM of each AuxRE was combined with 1, 2, or 5μg of either AtARF5-DBD or GmARF8a-DBD and subjected to SIPDI assay. An increase in fluorescence was observed with increasing amounts of ARF-DBD for both AuxRE variants, indicating that the amount of ARF (protein) was the limiting factor (Figure 4). These experiments also showed clear differences in binding strength between the concentrations of protein and DNA target, demonstrating that this assay has a high resolving power. Therefore, our approach of pulling down the protein-DNA complex using an affinity tag on the protein will be suitable to quantify relative strength of interactions efficiently.

**Figure 4.**
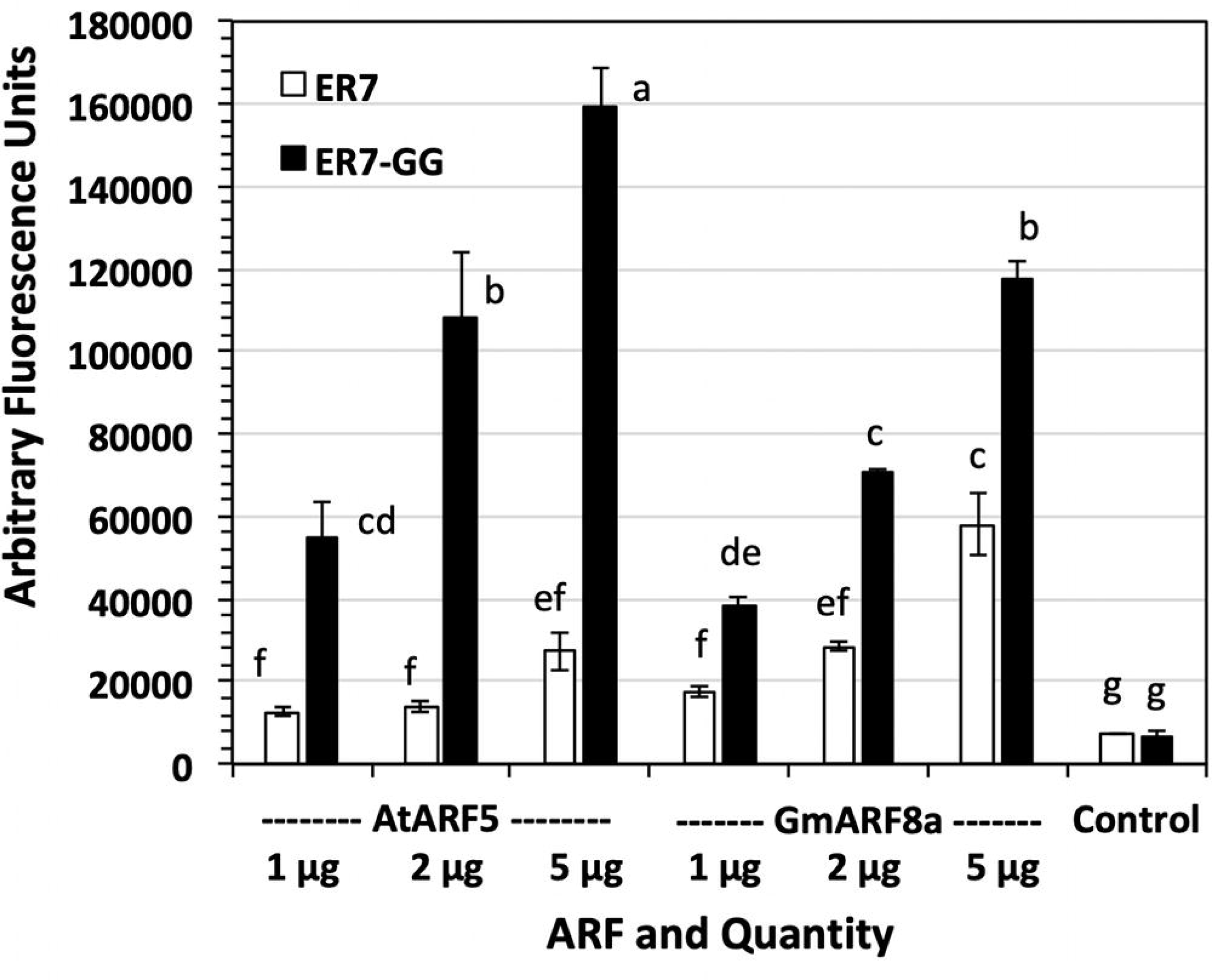
Binding of AtARF5 and GmARF8a to ER7 and ER7-GG when increasing amounts of protein were used in the assay. Reactions where no protein was added served as control. Data shown are average of three independent binding assays. Error bars indicate standard deviation. Fluorescence was measured using a BioTek Synergy 2 plate reader. Fluorescence measurements of samples labeled with distinct letters are significantly different from each other (ANOVA test, P < 0.05)

### Distinct affinities of two different activator ARFs to AuxRE variants

In the above experiment, we also compared the affinities of AtARF5-DBD and GmARF8a-DBD to the canonical ER7 motif (TGTCTC) and the ER7-GG motif (TGTCGG). Both ARFs showed much higher binding affinity to ER7-GG compared to ER7 as shown previously for the AtARF5-DBD (Figure 2; Boer, et al. (2014). Interestingly, while AtARF5 showed a 1.3-1.5 fold higher binding affinity than GmARF8a to ER7-GG (at 2 and 5 μg protein levels), GmARF8a-DBD showed about two-fold higher binding than AtARF5-DBD for ER7 (at 5 μg protein levels, Figure 4). These results suggest that different activator ARFs might have distinct binding strengths for AuxRE variants. This may contribute to a complex, yet fine-tuned auxin response.

These activator ARFs are also thought to preferentially bind as dimers as demonstrated by Boer et al (Boer, et al. 2014). An AuxRE where one of the TGTCTC motifs was changed to TGTCAA (ER7-1m) was used in the SIPDI assay to investigate the importance of ARF dimerization for proper DNA binding. Mutating one of the AuxRE motifs had a dramatic impact on the affinity of AtARF5 and GmARF8a, significantly reducing their binding. Interestingly, GmARF8a displayed significantly higher affinity than AtARF5 to ER7-1m, suggesting that even as a monomer 8a has a higher affinity for the TGTCTC motif (Figure 5). The reduction in affinity caused by the ER7 single motif mutation coincides with previous observations that the AuxRE facilitates ARF homodimerization and binding stability.

**Figure 5.**
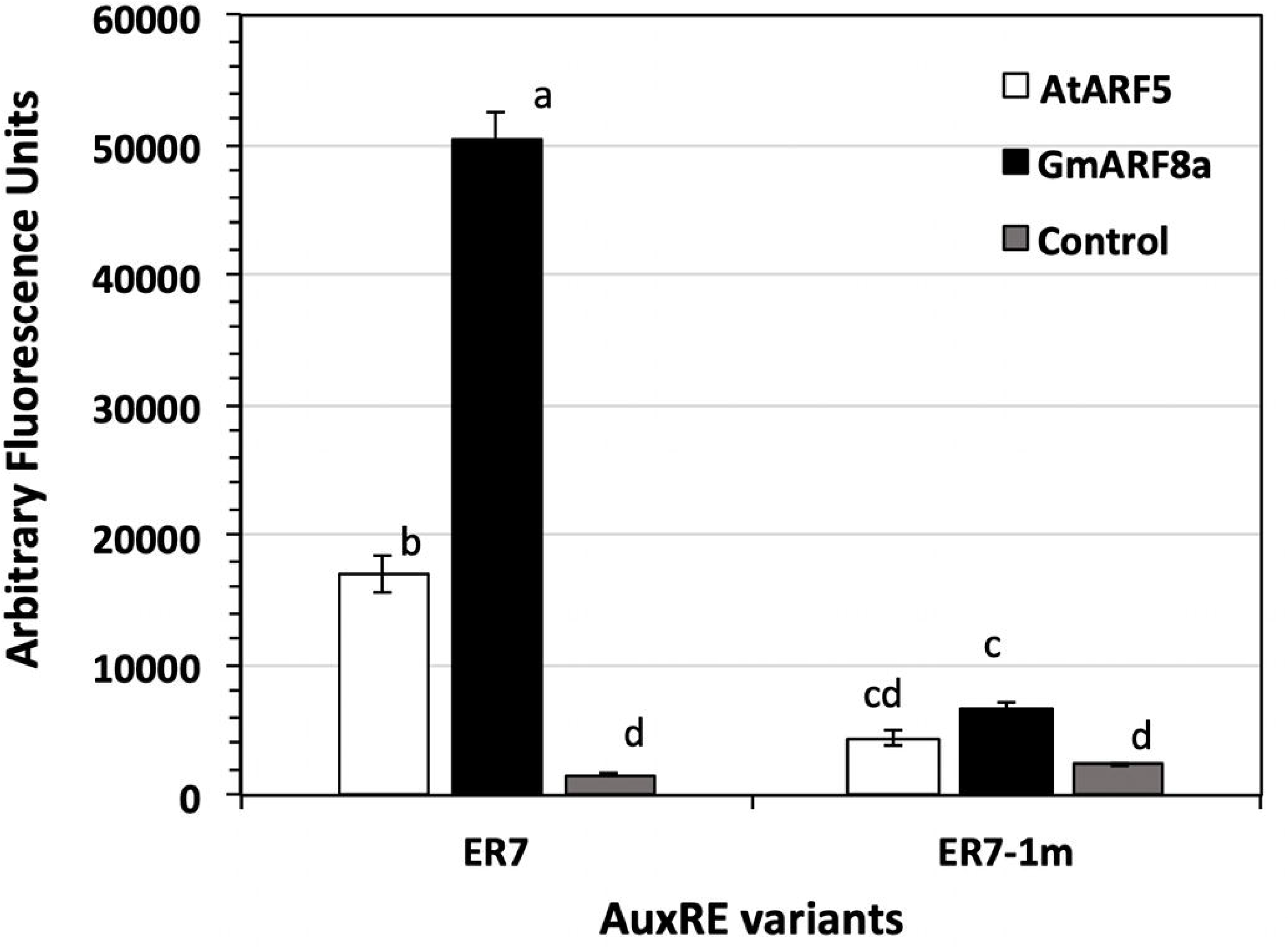
Binding of AtARF5 and GmARF8a to ER7 and ER7 mutated in one of the two binding sites (ER7-1m). Reactions where no protein was added served as control. Data shown are average of three independent binding assays. Error bars indicate standard deviation. Fluorescence measurements of samples labeled with distinct letters are significantly different from each other (ANOVA test, P < 0.05)

### SIPDI can be performed even with a UV gel doc system, making it more accessible

While the binding assay developed for this study used a plate reader to measure DNA binding as a function of fluorescence, its utility with more accessible instrumentation such as a gel doc was investigated. SYBR fluorescence in samples from a SIPDI assay performed using different amounts (1, 2, and 5μg) of each ARF or a no-protein control with saturating concentrations (6μM) of either ER7 or ER7-GG, were successively measured using a plate reader, LI-COR Odyssey imager, and UV transilluminator gel doc system. While the Odyssey imager and plate reader directly reported fluorescence measurements in arbitrary units, the gel doc image was further analyzed in ImageJ to obtain pixel intensity (see Experimental Procedures for details). The general trend of increased binding with increasing ARF concentrations was clearly observed in all three instruments (Figures 4, 6, and 7). However, the gel doc system (Figure 7) was not as efficient as the plate reader (Figure 4) or the LI-COR Odyssey (Figure 6) in distinguishing differences in binding at the lowest ARF concentration (1μg). This is likely due to the limited sensitivity of the platform compared to the other instruments. Nevertheless, there was at least 94% correlation between the gel doc and the other two sophisticated instruments (Table 1).

**Figure 6.**
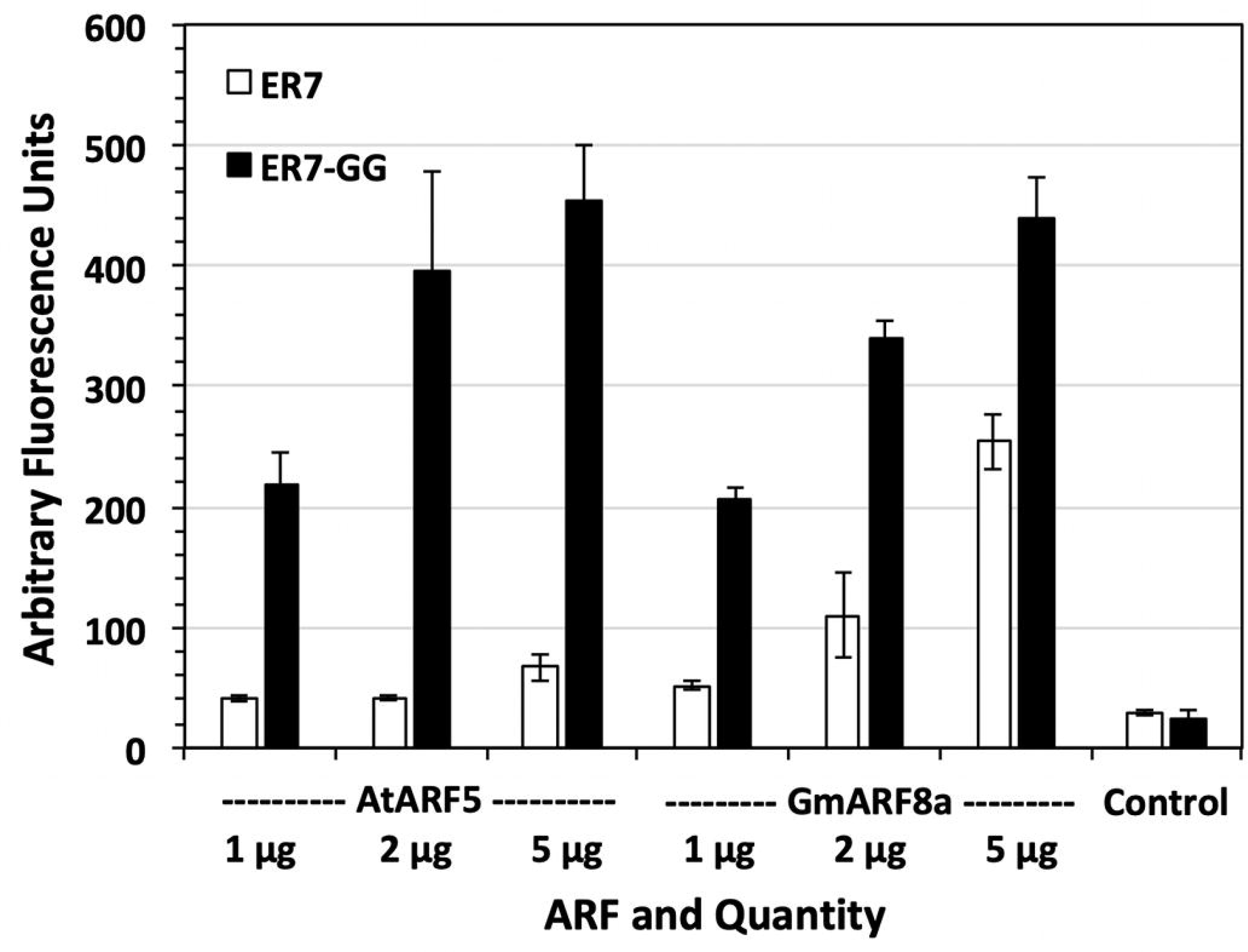
Binding of AtARF5 and GmARF8a to ER7 and ER7-GG when increasing amounts of protein were used in the assay. Reactions where no protein was added served as control. Fluorescence was measured using LI-COR Odyssey imager. Data shown are average of three independent binding assays. Error bars indicate standard deviation.

**Figure 7.**
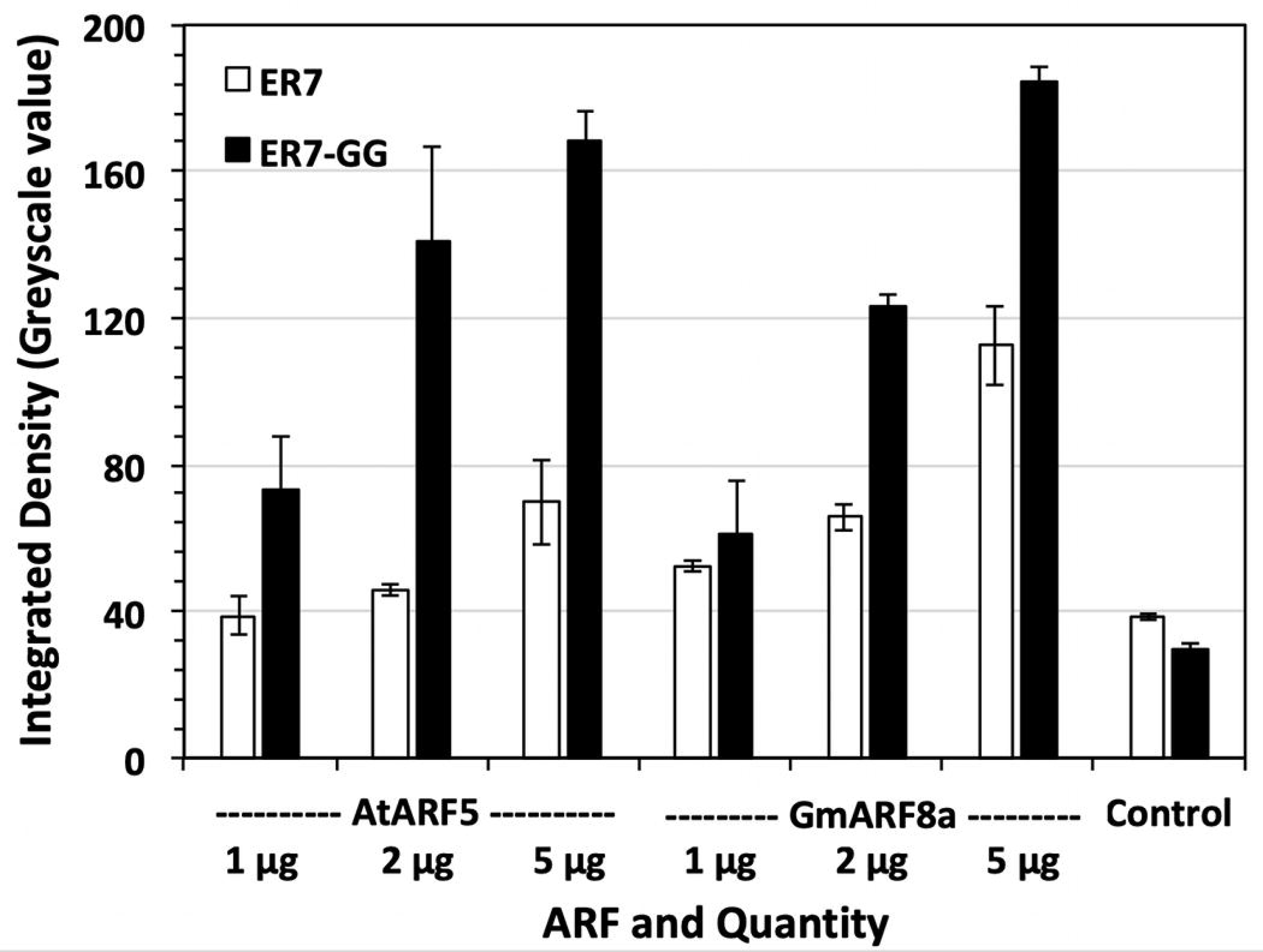
Binding of AtARF5 and GmARF8a to ER7 and ER7-GG when increasing amounts of protein were used in the assay. Reactions where no protein was added served as control. Fluorescence was measured by imaging on a UV Gel documentation system and subsequent image analysis using ImageJ. Data shown are average of three independent binding assays. Error bars indicate standard deviation.

**Table 1.** Correlation coefficients among protein-DNA binding values measured using a plate reader, LI-COR imager, and UV gel-doc system.

| *ER7* |  |  |  |
| --- | --- | --- | --- |
|  | *Plate reader* | *LI-COR* | *UV-box* |
| *Plate reader* | 1.000 |  |  |
| *LI-COR* | 0.976 | 1.000 |  |
| *UV-box* | 0.994 | 0.961 | 1.000 |
| *ER7-GG* |  |  |  |
|  | *Plate reader* | *LI-COR* | *UV-box* |
| *Plate reader* | 1.000 |  |  |
| *LI-COR* | 0.944 | 1.000 |  |
| *UV-box* | 0.937 | 0.971 | 1.000 |

The plate reader was more adept at discerning small differences at low fluorescence levels seen in the ARF quantities from 1 and 2 μg with ER7, the weaker-binding oligo based on one-way ANOVA comparisons performed separately on each oligo or each ARF. Therefore, it may be possible to extrapolate the amount of protein and DNA to determine approximate binding equilibrium. By comparison, it would likely be difficult to distinguish weaker-binding targets such as the ER7-single-mutation motif with low amounts of protein in the other platforms. However, 5μg of protein is certainly an attainable quantity and could supplement weak binding with the addition of more DNA target. Importantly, a highly accessible instrument such as a UV gel-doc, which is available in nearly all research laboratories and undergraduate institutions, can be used for the SIPDI assay for both research and educational purposes.

## Discussion

### Variation among ARF Affinities

In addition to the several molecular players that mediate auxin signaling, it appears that individual ARF binding strengths may also contribute to auxin’s fine-tuned activity in the cell. While both GmARF8a and AtARF5 had significantly higher binding strength than the no-protein controls, GmARF8a seems to interact more strongly with the AuxREs tested in this study. Notably, GmARF8a had much higher affinity for the classical ER7 motif than AtARF5 (Figure 4). Furthermore, GmARF8a has significantly higher binding strength for the mutated ER7 sequence that possesses only one TGTCTC motif, reinforcing its higher affinity for this motif. Both ARF DBDs seem equally capable of binding ER7-GG at saturating concentrations of protein and AuxRE target. However, at lower protein concentrations, AtARF5 seems to have stronger affinity for ER7-GG than GmARF8a (Figure 5). Galli *et al.* 2018 determined genome-wide binding sites of 14 maize ARFs using DAP-Seq and identified distinct binding sites of class A (activator) and class B (repressor) ARFs. Within the activator ARFs, the top binding motifs for ZmARF4 and ZmARF29, maize orthologs of AtARF5 were TGTCG and TGTCGG respectively. The study did not evaluate ZmARF3 and ZmARF30, the closest orthologs of GmARF8a. However, ZmARF34, the closest homolog of GmARF8a preferably bound TGTCT. These closely resemble our results suggesting that the binding preferences of ARF sub-clades might be evolutionarily conserved. These differences in binding affinity coupled with auxin concentration-dependent interaction between specific TIR/AFBs and Aux/IAAs (Calderon Villalobos *et al.* 2012, Shimizu-Mitao and Kakimoto 2014), can direct a precise but complex cellular response to auxin.

While there are affinity differences between the ARFs for the ER7 and ER7-GG motif, they expectedly had strong affinity for these sequences. Mutating one of the two TGTCTC motifs in the ER7 sequence nearly abolished affinity for AtARF5 and GmARF8a for this target, emphasizing that both AuxRE motifs coordinate dimer stability as previously demonstrated (Boer, et al. 2014). This also seems to highlight an evolutionary conservation within the ARF protein family as these ARFs are from different species. While not explored in this work, the optimum spacing between the AuxRE motifs may also be different between AtARF5 and GmARF8a, as well as their leniency for further mutations in one or both AuxRE motifs. Further investigations may examine how modifying single motifs of the AuxRE’s effect ARF binding (such as TGTCTC/TGTCGG), or how spacing between the motifs differs among these ARFs. Indeed, maize DAP-seq data suggested that motif orientation of and spacing between AuxRE sites are distinct between ARF clades (Galli, et al. 2018). The SIPDI assay could combine multiple components of the auxin signaling pathway (e.g. an ARF and Aux/IAA complex) to evaluate how these protein interactions influence ARF-DNA binding. There are certainly many possible sequence and/or spacing scenarios to explore, and this assay could provide a feasible solution for better understanding how the AuxRE collaborates with the auxin-responsive transcription factors to carry out auxin signaling.

### Utility of the assay for other applications

Banasik and Sachadyn have demonstrated the feasibility of Pierce nickel-coated plates as a useful tool for testing DNA-binding affinity (Banasik and Sachadyn 2016). However, these assays are not easily scalable due to the limited protein binding capacity in these plates. In addition, the plates did not permit efficient washing of unbound DNA leading to higher background fluorescence (unpublished observations). The SIPDI assay is scalable and provide minimal background as the amount of binding resin can be changed as needed and efficient washes are possible through centrifugation.

As anticipated, the plate reader produced sensitive results with little variation and is responsive to a large concentration range. The LI-COR Odyssey imager shows similar capacity for discerning differences between binding strength, but with a larger variation among the results. This variation could be problematic when investigating minute affinity differences. The UV transilluminator seems to be the least effective of the three, showing larger variation and is not as effective for parsing out small differences in binding strength like the plate reader. However, as instruments that are nearly ubiquitous in molecular biology laboratories (especially the UV transilluminator), gel doc systems are certainly useful for illuminating relative binding strength among proteins of interest and may be important for preliminary target screening prior to applying more advanced methods. Importantly, the assay performed using a gel doc system can be effective in teaching laboratories to demonstrate sequence-specific binding of proteins to DNA. Furthermore, the array of fluorophore technologies and the principle of this assay could likely be extended to testing other types of interactions, such as protein-protein (Chakravarty *et al.* 2015) or protein-RNA interactions. In addition, the assay can accommodate multiple molecular players during the initial binding step and help investigate how multi-protein interactions influence affinity for a specific target. Theoretically, this assay could also be used to uncover novel or unknown interaction partners to a protein or other molecule of interest. For example, a transcription factor whose binding sequence is not known could be mixed with a library of DNA sequences and applied to the assay. Interacting complexes that are maintained after the assay’s washes could then be cloned and sequenced by TA cloning or by PCR if the targets have flanking primer sequences.

In conclusion, we developed a simple yet reliable assay to quantify DNA-protein interactions in an accessible manner. Using the ARF-AuxRE interaction for validation, we determined that this assay can discern the relative strength of molecular interactions in a fast and accurate manner. We found there are preferential affinities between activator ARFs for AuxRE targets. In addition, the importance of dimerization within these ARF DBDs cannot be underscored and add another factor that contributes to successful ARF-DNA binding. These binding subtleties were examined on various platforms that ranged in sophistication. While the plate reader was most efficient at discerning the affinity differences, the Odyssey imager and UV transilluminator still provided useful data and confirmed the assay’s utility in virtually any laboratory setting including teaching labs in small colleges.

## Acknowledgements

The authors thank Prof. Dolf Weijers and his lab, Wageningen University, for providing the activator ARF coding sequences and helpful discussions concerning ARF activity. Research grant awards from USDA-NIFA-AFRI (2016-67014-24589), the National Science Foundation/EPSCoR Cooperative Agreement #IIA-1355423, and SD Agricultural Experiment Station (SD00H543-15) are gratefully acknowledged.

