## Supplementary Information for "A simple and sensitive SYBR Gold-based assay to quantify DNA-protein interactions"

***Supplementary data***

### Supplementary Tables

**Table S1.** Sequences of AuxRE variants used in the study. The differences last two nucleotides in TGTCTC motif or the spacer length between the evert repeat motifs are underlined.

| Target Name | Sequence |
|---|---|
| ER7 | 5′ CCGGTAGGT TGTCTC CCAAAGG GAGACA ACCGGTAGG<br>3′ GGCCATCCA ACAGAG GGTTTCC CTCTGT TGGCCATCC |
| ER7-GG | 5′ CCGGTAGGT TGTCGG CCAAAGG CCGACA ACCGGTAGG<br>3′ GGCCATCCA ACAGCC GGTTTCC GGCTGT TGGCCATCC |
| ER7-GA | 5′ CCGGTAGGT TGTCGA CCAAAGG TCGACA ACCGGTAGG<br>3′ GGCCATCCA ACAGCT GGTTTCC AGCTGT TGGCCATCC |
| ER7-AT | 5′ CCGGTAGGT TGTCAT CCAAAGG ATGACA ACCGGTAGG<br>3′ GGCCATCCA ACAGTA GGTTTCC TACTGT TGGCCATCC |
| ER5 | 5′ CCGGTAGGT TGTCTC CCAGG GAGACA ACCGGTAGG<br>3′ GGCCATCCA ACAGAG GGTCC CTCTGT TGGCCATCC |
| ER7-single motif | 5′ CCGGTAGGT TGTCAA CCAAAGG GAGACA ACCGGTAGG<br>3′ GGCCATCCA ACAGTT GGTTTCC CTCTGT TGGCCATCC |

**Table S2.** Sequences of primers used to clone the ARF DBDs.

| Primer Name | Sequence |
|---|---|
| pETite ARF8a DBD Fp | CGCGAACAGATTGGAGGTATGAAGCTTTCAACATCAGGGTTGG |
| pETite ARF8a DBD His Rp | GTGGCGGCCGCTCTATTAATGATGATGATGATGATGTAAAGGCTCAATTTCCCATAATGATACC |
| pETite ARF5 DBD Fp | CGCGAACAGATTGGAGGTATGATGGCTTCATTGTCTTGTGT |
| pETite ARF5 DBD His Rp | GTGGCGGCCGCTCTATTAGTGATGGTGGTGATGATGCGGTGTTTCGATATCCCATG |

**Table S3.** Buffers used in protein purification and SIPDI assay. All buffers were stored and used at 4ºC.

| Lysis Buffer | Wash Buffer | Elution Buffer (250mM) | Elution Buffer (500mM) | SPR Buffer |
|---|---|---|---|---|
| 50mM Sodium Phosphate Buffer | 50mM Sodium Phosphate Buffer | 50mM Sodium Phosphate Buffer | 50mM Sodium Phosphate Buffer | 20mM HEPES |
| 150mM NaCl | 150mM NaCl | 150mM NaCl | 150mM NaCl | 150mM NaCl |
| 40mM Imidazole | 40mM Imidazole | 250mM Imidazole | 500mM Imidazole | 1mM EDTA |
| 0.5mg/ml lysozyme (human) | | | | 0.01% Tween 20 |
| 0.01% IGEPAL | | | | 5 mM b-mercaptoethanol |
| | | | | *5M NaCl for salt treatment* |

### Supplementary Figures

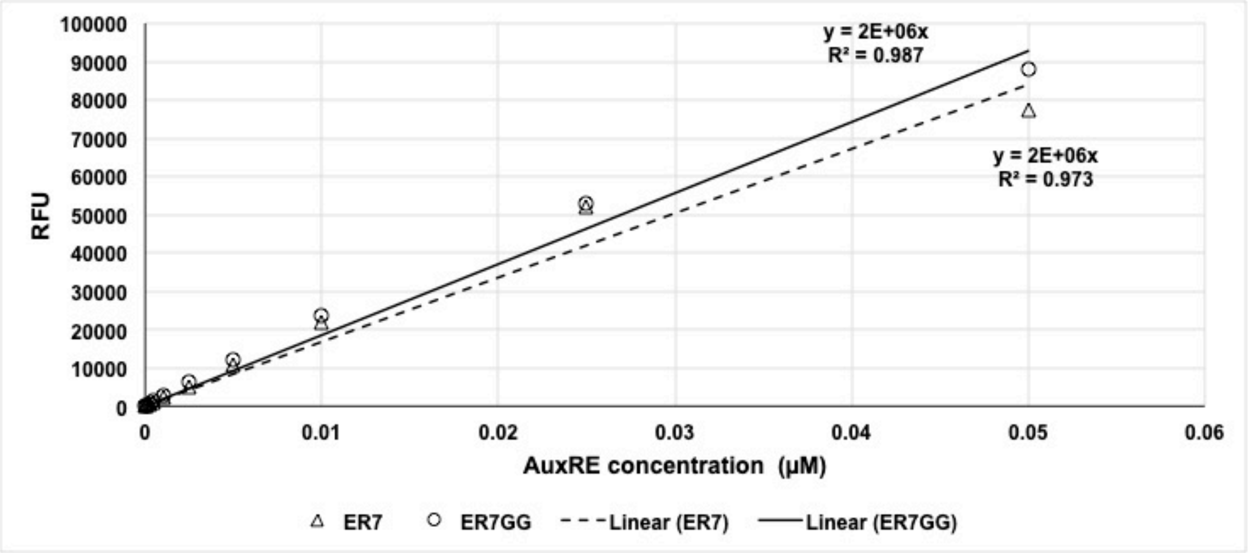


**Figure S1** Linear relationship between dsDNA concentration and SYBR Gold fluorescence assayed using a BioTek Synergy 2 plate reader.

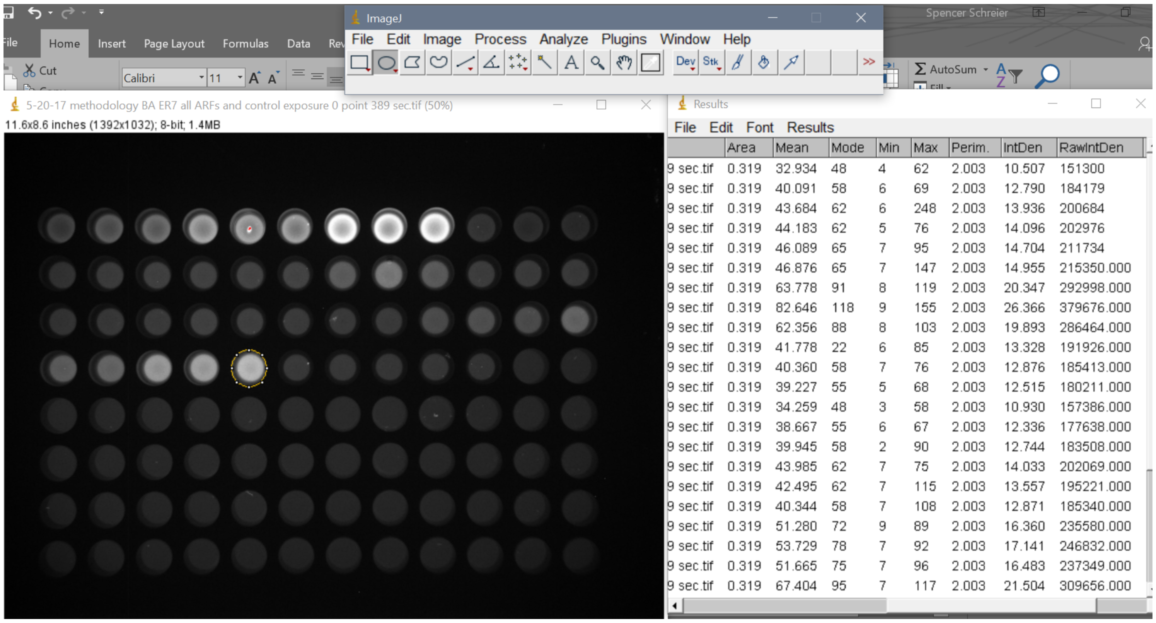


**Figure S2** A screen shot showing ImageJ-based quantification of SYBR Gold fluorescence on an image acquired using a UV Gel documentation system.

Image Display Values

| Channel | Color | Minimum | Maximum | K |
|---|---|---|---|---|
| 600 | Blue | 0.00119 | 0.598 | 0 |

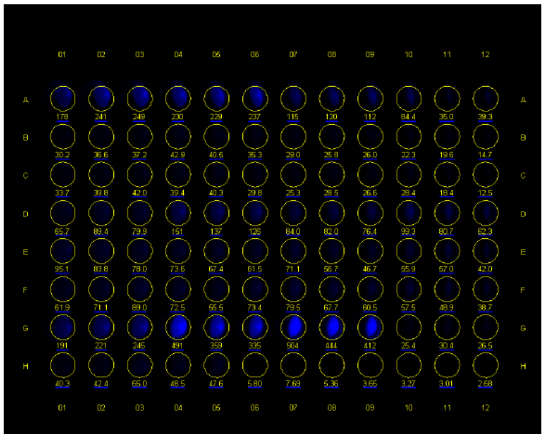

**Figure S3** Imaging and analysis parameters used to determine SYBR Gold fluorescence using a LiCor Odyssey imager. An image of a plate acquired using the parameters is shown as example.
